# When temporal attention lacks confidence

**DOI:** 10.1101/496232

**Authors:** Samuel Recht, Pascal Mamassian, Vincent de Gardelle

## Abstract

Accurate decision-making requires estimating the uncertainty of perceptual events. Temporal attention is known to enhance the selection of a stimulus at a relevant time, but how does this selective process affect a decision’s confidence? Here, we adapted an “Attentional blink” paradigm to investigate the effect of temporal attention on confidence judgments. In a RSVP stream of letters, two targets were cued to induce two successive attentional episodes. We found that the confidence ratings given to an item systematically followed the probability with which this item was reported. This coupling made confidence oblivious to selection delays usually observed when the two targets were separated by long intervals (249ms to 747ms). In particular, during this period, confidence was higher for more delayed item selection. One exception to this relationship between confidence and temporal selection was found when the second target appeared soon after (83ms) the first attentional episode. Here, a strong under-confidence bias was observed. Importantly, however, this early confidence bias did not impact confidence sensitivity in discriminating correct and erroneous responses. These results suggest that temporal attention and confidence can operate at different time scales, a difference which seems to reflect high-level heuristic biases rather than segregated processes for decision and confidence evidence.

## Introduction

Visual confidence is the subjective estimation of the accuracy of a decision made about a visual stimulus, and can be defined as self-evaluation of performance (Mamassian, 2016). It typically correlates with the objective accuracy of the decision, such that confidence can be used to regulate behavior and allocate resources when performance needs to be maintained or improved (Desender, Boldt, & Yeung, 2018; Guggenmos, Wilbertz, Hebart, & Sterzer, 2016; Hainguerlot, Vergnaud, & De Gardelle, 2018). However, humans do not always monitor their performance perfectly, and dissociations between confidence and performance have been documented in the literature (e.g., Graziano & Sigman, 2009; Lau & Passingham, 2006; Peters et al., 2017; Rahnev et al., 2011; Rahnev et al., 2012). Here, our goal is to assess how observers’ confidence judgments are affected by temporal attention, which is a major determinant of visual performance, even when stimulus’ characteristics are kept constant. In particular, we will rely on the “Attentional blink” phenomenon, by which participants can be blind to the second of two targets presented in close succession. The present work thus investigates how the temporal structure of the Attentional blink, which involves orienting attention towards two distinct moments, is tracked by confidence judgments.

Spatial attention enhances and prioritizes a stimulus representation at a particular point in space and inhibits other locations (Carrasco, 2011), while temporal attention enhances the selection and prioritization of a stimulus at a particular time (Coull & Nobre, 1998) and inhibits other time points (Denison, Heeger, & Carrasco, 2017). Both attention and confidence are related to discriminability: when attention increases the signal-to-noise ratio of an attended stimulus and facilitates discrimination, an efficient confidence estimate should, ideally, track this facilitation. Attention and confidence have already been considered in the spatial domain, leading to mixed findings. Some studies observed a dissociation between the two (Rahnev et al., 2011, 2012; Schoenherr, Leth-Steensen, & Petrusic, 2010; Wilimzig, Tsuchiya, Fahle, Einhäuser, & Koch, 2008), while other works tend to suggest that attention-dependent spatial uncertainty is usually well incorporated into confidence (Denison, Adler, Carrasco, & Ma, 2018; Zizlsperger, Sauvigny, & Haarmeier, 2012; Zizlsperger, Sauvigny, Händel, & Haarmeier, 2014). An open question relates to temporal attention and its link to confidence. This question is particularly relevant given the possibility that attention and confidence operates at different time scales (Rahnev, Koizumi, McCurdy, D’Esposito, & Lau, 2015), and that the temporal resolution of confidence during attentional orienting has not been fully investigated yet (Baranski & Petrusic, 1994).

In some circumstances, temporal attention can be suppressed, delayed or misplaced. One relevant and very robust finding regarding this issue is the Attentional blink phenomenon (Broadbent & Broadbent, 1987; Raymond, Shapiro, & Arnell, 1992). Specifically, when two targets are embedded in a rapid serial visual presentation stream, the second target T2 is often missed when it appears soon after (150-300ms later) the first target T1. This phenomenon of missing the second target is called “Attentional blink”. When temporal selection of T2 is not simply suppressed (in the case of missed T2 targets), it may still be delayed, such that an item following T2 would be reported instead of T2. These selection delays, sometimes known as “post-target error intrusions” (Chun, 1997; Vul, Hanus, & Kanwisher, 2008) are a second feature of the Attentional blink. Finally, when T2 is presented immediately after T1 (60-100ms), then both targets are on average accurately reported. This special case has been coined “lag-1 sparing” (e.g., Hommel & Akyürek, 2005) and is a third feature of the Attentional blink. All three features of the phenomenon can be accounted for by a variety of models, some based on the idea of resources-depletion and others built from fundamental principles of selection (for reviews, see Dux & Marois, 2009; Martens & Wyble, 2010). However, whether confidence tracks these features remains an open empirical question. To address this question in the present study, we used an Attentional blink paradigm in combination with confidence judgments for both T1 and T2 reports.

The empirical question considered here is whether participants’ confidence would reflect these three basic features of the Attentional blink, namely (1) the drop of performance, (2) the delayed temporal selection that follows, and (3) the lag-1 sparing effect. To measure errors and delays in temporal selection, one can present participants with a rapid stream of letters, and indicate two letters in the stream for latter report. The serial position of each letter in the stream provided critical information on the point in time at which attention was deployed (Goodbourn et al., 2016; Martini, 2012; Vul, Nieuwenstein, & Kanwisher, 2008). In other words, the present work proposes to investigate whether participants accurately evaluate the limits of their ability to deploy their attention at the right moment in time.

## Material and methods

### Participants

39 adult volunteers were recruited from the Laboratoire d’Economie Expérimentale de Paris (LEEP) pool of participants (M ± SD = 25.5 ± 2.9 years old). They all provided informed written consent prior to the experiment. Four observers were discarded following a technical problem, and three participants were removed following extremely small accuracy rate for target 1 or 2 (exclusion criterion: <10% accuracy), leaving 32 participants for analysis. Observers were paid a base sum (10 EUR) plus a bonus depending on their performance in the task (up to 10 EUR in addition). The average payoff was 16.43 EUR (± 1.89) for a single 1.5 hours session. The experimental procedure received approval from the Paris School of Economics (PSE) ethics review board and adhered to the principles of the Declaration of Helsinki.

### Apparatus and stimuli

Participants sat approximately 60 cm from the screen (1280×1024 pixels, 60 Hz refresh rate). Stimuli were generated using the Python programming language and the PsychoPy toolbox (Peirce, 2007) on a Windows XP computer. On each trial, participants were presented with a rapid serial visual presentation (RSVP) stream of the 26 English letters (Courier New, white font, 2.5° of visual angle) in the center of a black screen background (Figure 1). Letters were randomized, and each letter was presented for 33ms (3 frames) with an inter-stimulus interval of 50ms (4 frames), which corresponds roughly to a 12 Hz stream. Two letters in the stream were targets surrounded by a visual cue (white annulus, inner/outer diameter: 2.9°/3.1°), which appeared simultaneously with the target. The first target (T1) was located between the 5th and the 10th item in the stream, while the second target occurred at the 1st, 2nd, 3rd, 6th or 9th position after T1. Both target positions where counterbalanced with a full factorial design.

**Figure 1.**
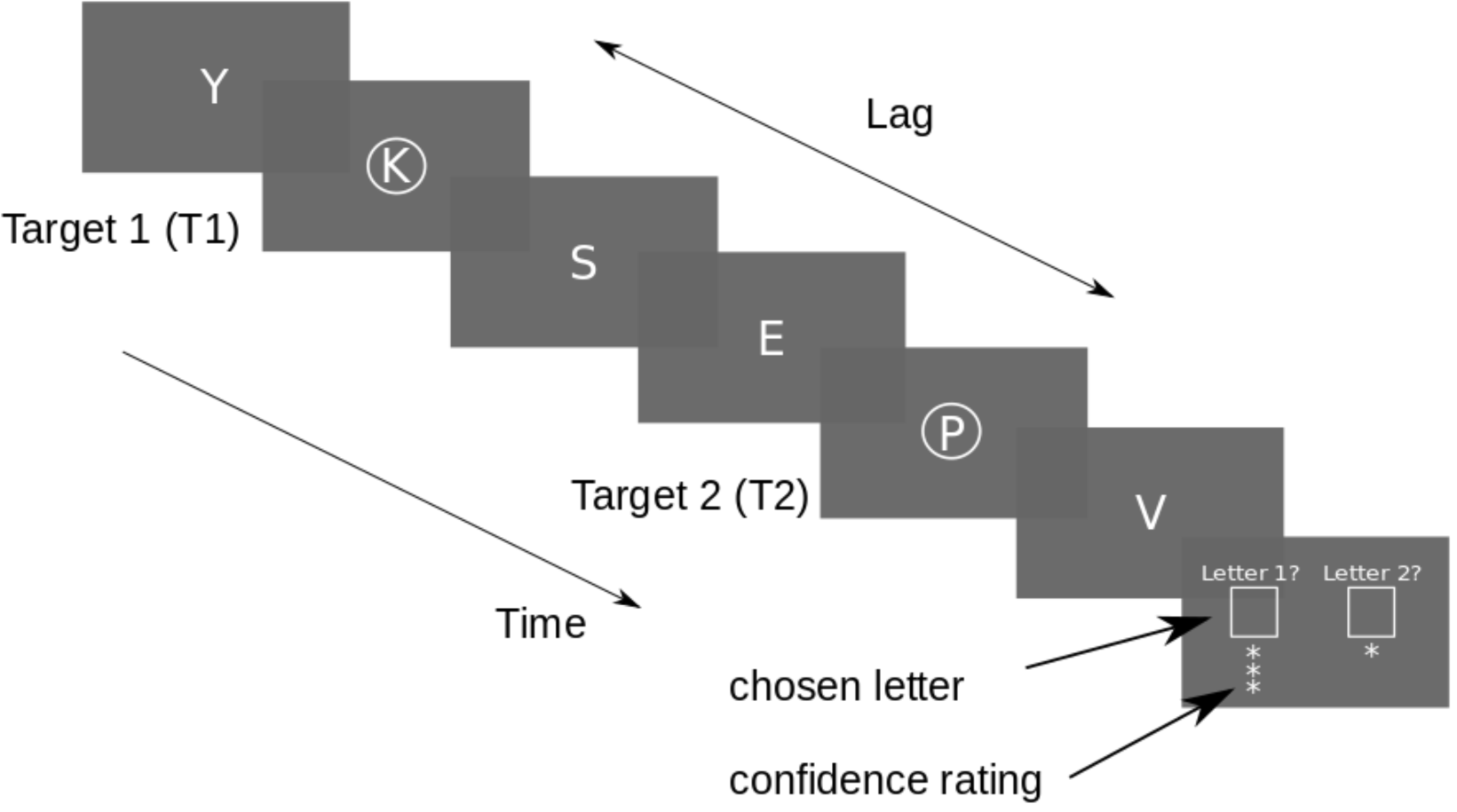
Experiment design: participants were required to report the two cued letters in the RSVP, and rate their confidence for each reported letter (Experiment 1) or for only one of the letter (Experiment 2, see Supplementary material) on a three-point scale. The distance in items (or lag) between the first and second target was varied (lag-3 depicted here). Each letter appeared for 33ms, followed by a 50ms ISI.

The lags between T1 and T2 were chosen so as to sample the different periods of the Attentional blink: lag-1 (83ms after T1), where lag-1 sparing is known to occur; lag-2 and 3 (166ms and 249ms), which usually show strong drop in T2 reporting accuracy; and finally lags 6 through 9 that demonstrate a progressive recovery in accuracy (498ms and 747ms).

### Procedure

At the end of each trial, participants had to report each target letter, in order of appearance, as well as their confidence for each report. Duplicates of the same letter were not accepted, given that each letter only appeared once in the stream. Confidence was rated using a three-point scale and incentivized. Specifically, participants were informed that each of their responses would generate a number of points equal to their confidence (with points worth 0.5 EUR for correct responses and 0 EUR for incorrect responses), and that every 25 trials one point would be randomly selected from those generated in the past 25 trials. This scheme was applied separately to T1 and T2 responses. High accuracy and good confidence estimates were therefore decisive to maximize payoff. Participants did not receive accuracy feedback until the very end of the experiment.

Before the main experiment, participants completed 10 practice trials, the first half without confidence judgments. The main session then consisted in 500 trials, with a 10-seconds break every 6 minutes.

## Analyses

All the analyses were carried out using the R programming language. Mixed effects models were built using the Lme4 R package. Accuracy and average confidence of T1 and T2 reports were analyzed using standard ANOVAs. In the current paradigm, the position of the reported item is also of interest. To analyze how reports and confidence depended on this serial position, a mixed effects model comparison approach was used. Specifically, a regression with fixed effects of position (and possibly other factors) and participants as random intercepts was compared to a regression without the fixed effect of position. When necessary, a third model including an interaction was added to the comparison.

Statistical results involving serial positions were systematically confirmed using permutation analysis, given the unbalanced nature of the dataset in this case. Serial positions were randomly shuffled for each participant and lag separately (for the whole dataset) and the relevant statistical analysis was applied to this surrogate data. The process was repeated 3,000 times, and the resulting distribution was compared to the test result obtained on the original data. P-values obtained through this method are reported as ‘p_RAND_’.

When necessary, ANOVAs were corrected using the Greenhouse-Geisser adjustment and t-tests were corrected using the Welch-Satterthwaite adjustment. We report Wilcoxon signed ranked test using uppercase T when the Shapiro-Wilk normality test failed, and Student test using lowercase t otherwise.

## Results

### Overview

We started by analyzing T1 data to obtain a baseline in which attention is not challenged. We evaluated how T1 reports were distributed around the target’s true position, and how confidence differed between correct reports and errors. We further evaluated how confidence varied within errors, as a function of the position of the reported item relative to the target. Focusing on T2, and the classical Attentional blink phenomenon, we evaluated the variation of both performance and confidence across lags. There are three main findings. First, confidence captures the Attentional blink but elicits a strong dissociation from accuracy during lag-1 sparing. Second, confidence was oblivious to the delays in item selection induced by the Attentional blink. Third, confidence generally followed the likelihood of selecting items, irrespectively of whether the selection is targeted at T1 or T2, correct or incorrect, or delayed by the Attentional blink.

### T1: distribution of reports and confidence

The task of reporting the first target is considered here as a near-baseline temporal selection process during initial orienting of attention. Overall, T1 targets were identified correctly 43% of the time. As can be seen on Figure 2A, and as documented previously (Vul, Nieuwenstein, et al., 2008), errors were not random guesses. The letter presented just before or just after the target was reported in 18% of the trials, largely exceeding the guess rate of 1/26≈4% (t(31)=21.7 p<0.001). Focusing on serial positions from 2 items before to 2 items after T1 (included), we further tested how report frequency can be predicted from the position, the lag and their interaction (using mixed models, see Analyses). Including item position as a predictor outperformed a model without the position effect (χ^2^(4)=1058, p_RAND_<0.01). Including the lag x position interaction improved the model even further (χ^2^(16)=43.3, p_RAND_<0.01), but this interaction seemed specifically driven by the lag-1 as it disappeared when excluding this lag from the analysis (χ^2^(16)=5.6, p_RAND_=0.95). The interaction between lag and position might reflect the confusion and order reversals that occur at lag-1 (see Supplementary material).

**Figure 2.**
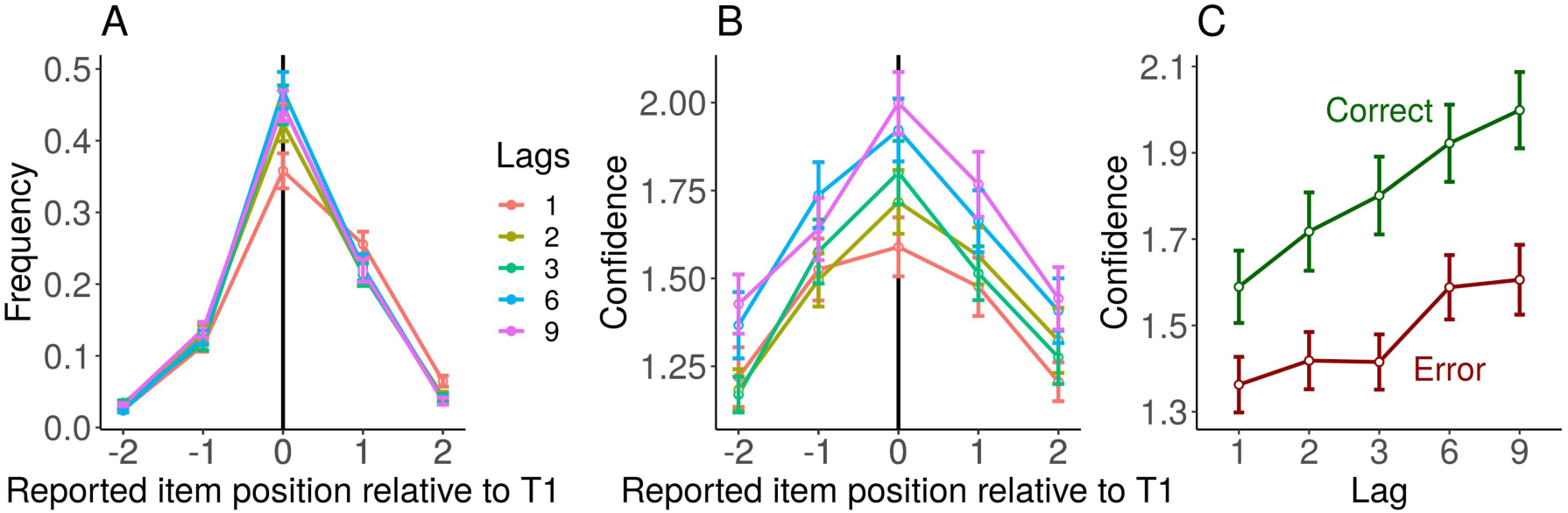
Reports and confidence about initial attentional orienting (T1): (A) The frequency of report for item around target true position. (B) The corresponding average confidence per position. (C) The correct and error trials average confidence level, which provides an estimate of objective, error-based metacognition. Error bars represent standard error of the mean across participants.

One striking feature of the data is that confidence followed a profile similar to report frequency: when a specific position was reported more frequently, these reports were also associated with greater confidence (Figure 2B). Confidence was significantly affected by item position (χ^2^(4)=240, p_RAND_<0.01). Including the interaction between lag and position however did not improved the model (χ^2^(16)=15.8, p_RAND_=0.55). We replicated these analyses while excluding correct responses, to confirm that these results did not merely reflect the ability to discriminate between correct and erroneous responses (see “error-based metacognition” below).

To directly evaluate the similarity between confidence and report frequency, confidence was averaged for each participant by grouping all lags together, and we correlated this average confidence to the report frequency, across the 5 report positions centred on the target (including the target’s position). The mean r coefficient was 0.86, across participants (95% CI=[0.82 0.90]; t(31)=44.2, p_RAND_<0.001). Thus, it appears that participants’ confidence is closely linked to the probability with which the reported letter is selected.

### T1: Error-based metacognition

One typical signature of metacognitive ability is the difference of confidence between correct and incorrect reports, with higher confidence for correct responses. Figure 2C illustrates this measure for the different lags. A repeated-measures ANOVA with lag and trial type (correct vs. error) revealed a main effect of trial type (F(1,31)=77.8, MSE=0.11, p<0.001), a main effect of lag (F(2.04,63.4)=38.2, MSE=0.06, p<0.001), as well as a lag x type interaction (F(3.35,104)=5.7, MSE=0.02, p<0.001). Overall, participants gave higher confidence to correct than to incorrect T1 responses. This difference between trial types increased with the lag between T1 and T2, but was present for all lags (all p<0.01).

### T2: confidence tracks the Attentional blink but not Lag-1 sparing

We then turned on reports and confidence judgment about T2 targets (see Figure 3A), which corresponds to the initiation of a second attentional episode (or “reorienting” of attention in time). To make sure of a successful initial attentional capture by T1, the following analysis was restricted to trials in which T1 was correctly reported. In these trials, 23% of T2 reports were correct. Figure 3A shows T2 accuracy and confidence for the different T1-T2 lags. T2 accuracy was affected by the lag between T1 and T2 (F(2.14,66.5)=67.2, MSE=0.02, p<0.001) and exhibited the classical Attentional blink effect: it dropped for lag-2 and lag-3 relative to longer lags (2-3 vs. 6-9: T(31)=0, p<0.001).

**Figure 3.**
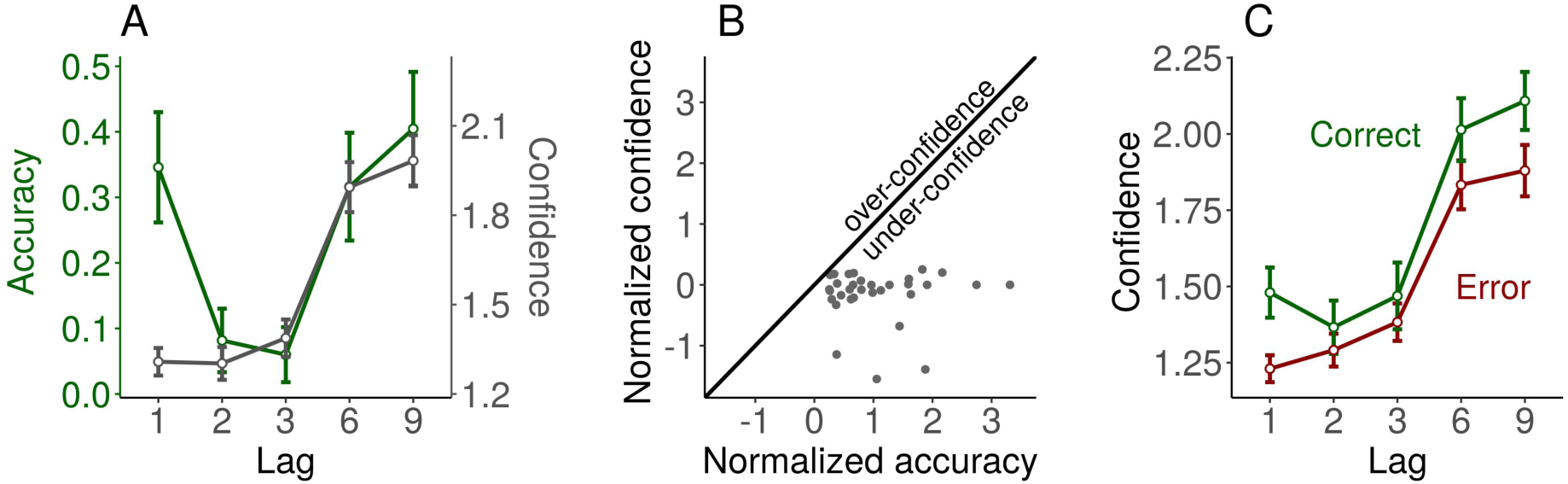
Attentional blink and early confidence bias: (A) T2 average accuracy (in green) and confidence (in grey) as a function of the lag between T1 and T2. (B) The systematic under-confidence occurring when attention reorients to a second point in time 83ms after the first target (Lag-1), by representing the accuracy and confidence of lag-1 as a point in the space inferred from the covariation of confidence and accuracy from lag-3 to lag-9. Each point is a participant. (C) The average confidence level for correct T2 reports and errors (error-based metacognition), for each lag. Metacognitive sensitivity is conserved at lag-1 despite a bias for low confidence ratings. Error bars represent standard error of the mean across participants.

**Figure 4.**
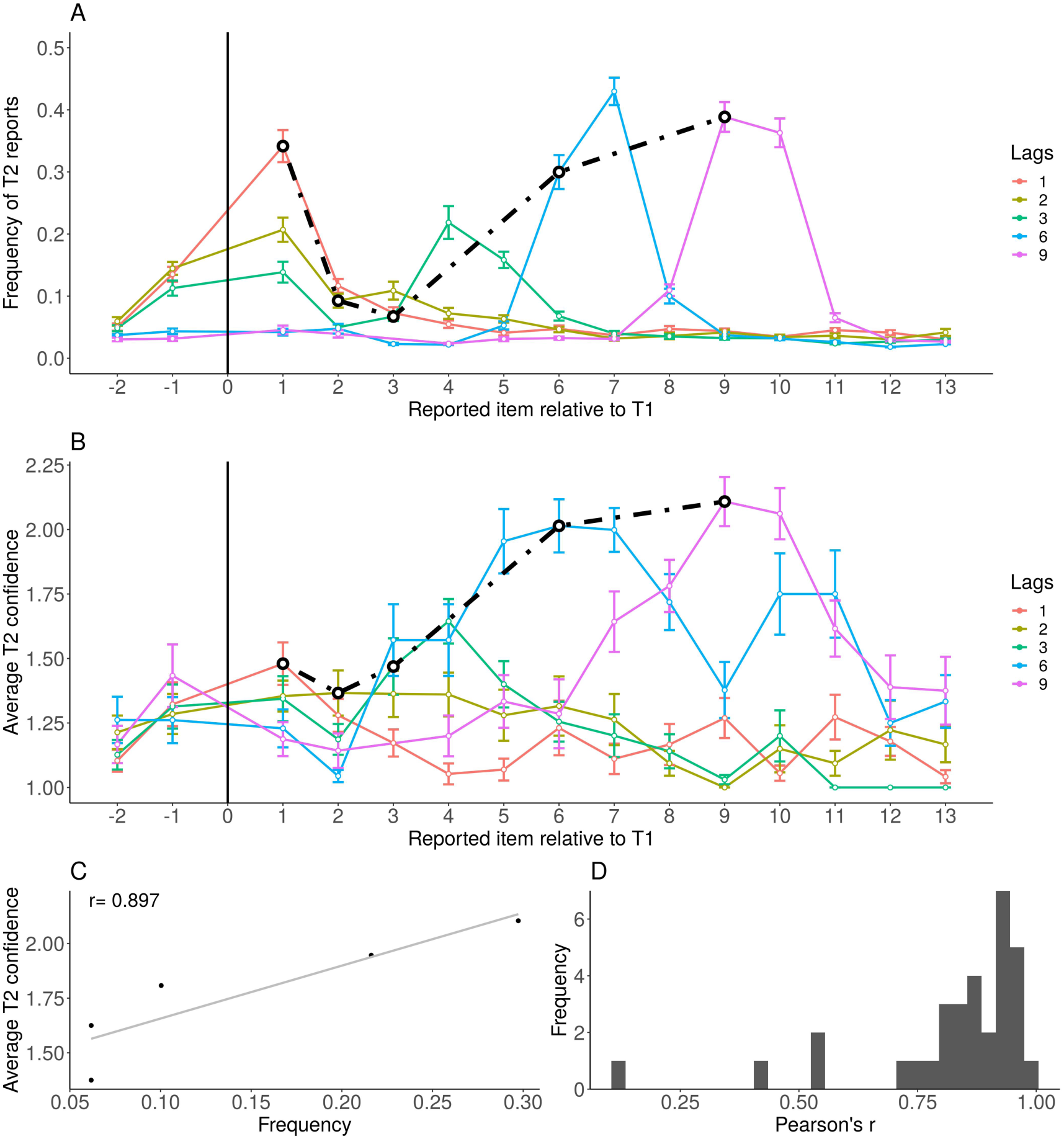
Reports and confidence about attentional reorienting (T2): (A) The frequency of T2 reports as a function of the position of the reported item relative to T1, for each lag. Note that T1 position has no value, given that only trials in which T1 is correctly reported were considered here (hence T2 reports cannot correspond to T1 position). The black line connects the points corresponding to accurate T2 reports. (B) Confidence of the T2 reports, as a function of the position of the reported item relative to T1, for each lag. The black line connects the points corresponding to accurate T2 reports. Error bars represent standard error of the mean across participants. (C) Regression between frequency and confidence with 5 positions centered on T2, collapsed across lags, for a representative participant. (D) Histogram of the correlation coefficients for all the participants. The confidence-frequency relation is strong and holds for most participants. Please refer to the online version of the paper for colors.

Confidence was also affected by lag (F(1.88,58.4)=92.4, MSE=0.08, p<0.001) and dropped for lags 2-3 relative to longer lags (2-3 vs. 6-9: T(31)=0, p<0.001), paralleling accuracy. Thus, participants were able to acknowledge the drop of performance at lags 2-3 relative to longer lags.

Importantly however, participants’ confidence was strongly dissociated from accuracy at lag-1. Confidence seemed blind to lag-1 sparing, a classical phenomenon where T2 accuracy at lag-1 is much higher than during the blink period (1 vs. 2-3: T(31)=528, p<0.001) and indistinguishable from long lags (1 vs. 6-9: T(31)=260, p=0.95). Indeed, lag-1 confidence was as low as for lag 2-3 (T(31)=197, p=0.66) and much lower than for long lags (1 vs. 6-9: T(31)=0, p<0.001). To further quantify this “lag-1 under-confidence”, we asked whether the increase in accuracy at lag-1 relative to lag-3 was accompanied by the expected increase in confidence, estimated from the covariation between accuracy and confidence between lag-3 and lag-9. To do so, for each participant we normalized the confidence at lag-1 with respect to its variation between lag-3 and lag-9 (i.e. across the Attentional blink), by applying the transformation x_1_ -> x_1_′ = (x_1_-x_3_)/(x_9_-x_3_) where x_k_ is the confidence at lag-k. We applied the same transformation to accuracy. Figure 3B shows the ‘normalized’ confidence and accuracy at lag-1, in this lag-3-to-9 space, where lag-3 and lag-9 have (0,0) and (1,1) coordinates, respectively. All participants are located below the diagonal, suggesting that they are less confident at lag-1 than what would be expected given their accuracy level at lag-1. This lag-1 under-confidence, computed as the average difference between predicted and observed lag-1 confidence, was significant at the group level (M=0.627, 95% CI=[0.448 0.807]; t(31)=7.1439, p<0.001).

### T2: Error-based metacognition

We then focused on error-based metacognition, defined above as the difference in confidence between correct reports and errors. Because some participants had no correct answers during the Attentional blink, only a subset of participants was considered here (N=24). As can be seen from Figure 3C, participants overall expressed higher confidence when they were correct (F(1,23)=11, MSE=0.15, p=0.002). This error-based metacognition interacted with the lag (F(3.46,83.1)= 3.28 MSE=0.05, p=0.02). Post-hoc Bonferroni-corrected tests showed that the difference in confidence between correct T2 reports and errors was significant for lag-1 (t(23)=3.7, p<0.01), lag-6 (t(23)=3.1, p=0.02) and lag-9 (t(23)=4.3, p=0.001) but not for lag-2 (t(23)=1.4, p=0.895) and lag-3 (t(23)=0.1, p=0.99). In other words, the ability to detect objective errors was diminished specifically during the Attentional blink period, and it did not disappear at lag-1, despite the low level of confidence.

### T2: Delays in temporal selection and confidence

To analyze the delay in selection and confidence following reorienting of attention to T2, we computed the average position of the reported item relative to the target position, in an 11-items window centered on the target position. This measure, called the “center of mass” (Goodbourn et al., 2016; Vul, Hanus, et al., 2008; Vul, Nieuwenstein, et al., 2008) is positive when a delay occurs in item selection. Figure 5B illustrates the distribution of the center of mass across participants, separately for each lag and for the 3 confidence levels, and shows that T2 item selection is delayed specifically after the Attentional blink (at lags 3, 6 and 9). A model comparison approach confirmed that including the lag as a predictor for the center of mass significantly outperformed the null model (χ^2^(4)=56.9, p_RAND_<0.001). Furthermore, a model including confidence, lag and their interaction as fixed effects outperformed a model without the interaction term (χ^2^(8)=30.1, p_RAND_<0.01), which in turn outperformed a model without the main effects of confidence and lag (χ^2^(2)=10.7, p_RAND_<0.01). When looking at each lag separately, we found that the effect of confidence on the center of mass was only significant for lag-6 (χ^2^(2)=9.2, p_RAND_<0.05) but not for the other lags after Bonferroni correction (p_RAND_>0.1). At lag-6, high confidence reports are associated with longer delays than low confidence reports (Figure 5, right panel), confirming that confidence is oblivious to the delays induced by the Attentional blink. For comparison, similar analysis on T1 showed that longer delays are associated with lower confidence overall (Figure 5A). This analysis on T1 confirmed a significant effect of lag on the center of mass for both selection (χ^2^(4)=19.4, p_RAND_<0.001) and confidence (χ^2^(8)=9.5, p_RAND_=0.01), but no interaction for confidence (χ^2^(8)=6.5, p_RAND_=0.56).

**Figure 5.**
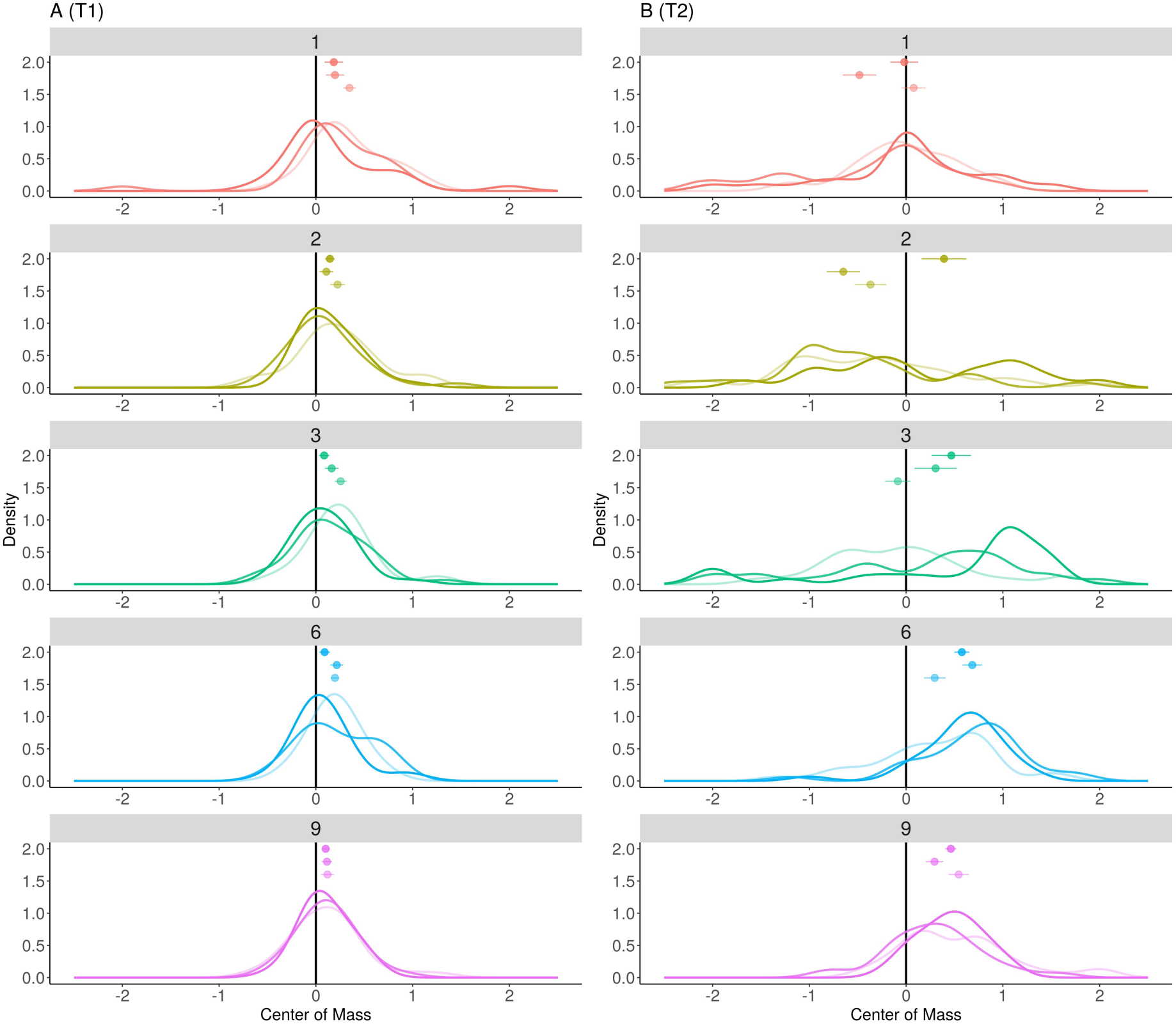
Center of mass analyses. Each panel represents for one lag the distribution of the center of mass across participants, relative to the T1 or T2 positions. The 3 confidence levels are plotted separately for each lag, with more transparent lines for lower confidence. Note that at lag-6, higher T2 confidence is associated with longer delays, suggesting that confidence does not accurately track the delay of temporal attention. Dots and error bars represent the mean and standard error of the mean for each distribution. Densities were estimated using Gaussian kernels (sd: 0.2).

### T2: Confidence follows report frequency

To evaluate more precisely the mediator of confidence delay following the Attentional blink, we considered - for the different lags - the positions of the reported items relative to the target (Figure 4A). As for T1, errors were not random guesses but distributed around the target, and items appearing just before or just after the target were reported more often (17%, with a 95% CI=[0.16 0.18]; vs. chance level at 4%: t(31)=19.5, p<0.001). At lag-6, the observed delay in selection and confidence described previously suggests that the most common error was to report the item presented immediately after the target, and rate it with higher confidence than correct reports.

Another way to look at the effect of position on confidence is to test whether high confidence ratings are assigned for positions reported frequently, as was found for T1 reports. Comparing Figures 4A and 4B, we note that for each lag confidence and report frequency typically peak at the same item position, even when this item position is not the target position. This similarity between confidence and report frequency was confirmed by looking at their correlation across lags for 5 positions centered on T2 (Mean r coefficient: 0.82, 95% CI=[0.76 0.89]; t(31)=25.3, p_RAND_<0.001), as shown on Figure 4D. Figure 4C plots the regression on one representative participant for illustrative purpose.

### A replication with a reduced metacognitive load (Experiment 2)

In Experiment 1, participants reported their confidence for both T1 and T2 targets in each trial. The high demand put on the metacognitive system during the task might explain why confidence failed to track the lag-1 sparing or the delays in item selection induced by the Attentional blink. To address this possibility, we conducted a second experiment in which we lowered the demands put on the metacognitive system, by asking only one confidence estimate per trial. In experiment 2, participants (N=29) gave their confidence about T1 in the first half of the experiment and their confidence about T2 in the other half (or vice-versa, counterbalanced across participants). All other parameters were identical to Experiment 1, and performance levels in Experiment 2 were similar to Experiment 1, with an average accuracy at 40% for T1 and at 22% for T2 after a correct T1 response.

Critically, in Experiment 2 we replicated the three main findings of Experiment 1, as summarized below (for details see the Supplementary material). First, participants were oblivious to lag-1 sparing and exhibited a clear under-confidence at lag-1 for their T2 reports (see Figure S2). Second, temporal selection was delayed after the Attentional blink but confidence judgments did not track this delay (Figure S3, A and B). Instead, the center of mass of T2 responses was more delayed when participants expressed greater confidence (Figure S4B), as was found in Experiment 1. In other words, whereas the metacognitive task was less demanding, participants were not better at acknowledging the lag-1 sparing or delays in temporal selection induced by the Attentional blink. Finally, we replicated the finding that confidence was tied to report frequency for T1 (Figure S1). Hence, when a particular item was more likely to be selected, it was also reported with a greater confidence.

## Discussion

The present study considered how human observers could evaluate their own performance in a task in which temporal attention reoriented after a variable delay following an initial attentional episode. To do so, confidence judgments were introduced within an Attentional blink paradigm, and we analyzed how such judgments would track the limits of performance typically observed in this experimental protocol. We obtained three main results. First, participants noticed the drop of accuracy for a second target (T2) caused by the Attentional blink but they failed to notice the sparing of T2 accuracy at lag-1 (83ms after the peak of the first attentional episode). Second, participants were oblivious to the delays in temporal selection induced by the Attentional blink. Third, participants’ confidence judgment about a selected item followed the likelihood with which this item would be selected in the sequence. All these results were replicated in a second experiment in which we only collected one confidence judgment (either on T1 or on T2), to reduce the demands put on the metacognitive system.

### Confidence during the Attentional blink and its early dissociation from accuracy

During the Attentional blink, confidence captured the drop in accuracy during lag-2 and lag-3, and the progressive recovery for longer lags (Figures 3A and S2A). Surprisingly, confidence was however not able to track the sparing of accuracy for the lag-1 (83ms after T1). This under-confidence at lag-1 was very clear, and present for almost every participant in two experiments (Figures 3B and S2B). Thus, confidence seems to only partially mirror visibility, which is degraded when the Attentional blink occurs, although mixed results were found for lag-1 in the literature (Nieuwenhuis & de Kleijn, 2011; Pincham, Bowman, & Szucs, 2016; Sergent & Dehaene, 2004; Simione, Akyürek, Vastola, Raffone, & Bowman, 2017). This similar pattern was not to be taken for granted, as confidence and visibility do not always go hand in hand, and can be dissociated both conceptually and empirically (Rosenthal, 2018).

However, while participants were clearly under-confident at lag-1, they were still sensitive at the metacognitive level, as they could discriminate between correct responses and errors, and between different errors. This under-confidence bias at lag-1 might result from participants being worried about the possibility of an order reversal (see Supplementary material), where T1 would be reported as T2 and vice-versa. This order-reversal phenomenon is well-known in the literature, and it has been suggested that at lag-1, T2 would actually benefit from T1 attentional episode, the two targets being often perceived as a single object (Akyürek et al., 2012; Goodbourn et al., 2016; Hommel & Akyürek, 2005), at the cost of an increased uncertainty about their relative order.

To summarize, when one attentional episode encompasses two responses, a confidence cost is applied to penalize the second reported item possibly because of increased risk of confusion between the two targets. This confusability risk could also induce a confidence bias favoring items further away from T1 attentional episode, which would result in delayed selection and confidence for T2.

### Delay in selection and confidence

Our second result relates to the delayed attentional selection induced by the Attentional blink (Chun, 1997; Vul, Hanus, et al., 2008). We found in both experiments a maximal delay at Lag-3, which confidence ratings could not capture. On the contrary, at this lag, we found higher confidence when item selection was more delayed, a tendency that was significant at lag-6 (Figure 4 and Figure 5) for Experiment 1 and at lag-3 for Experiment 2. The similarities between delays in attentional selection and confidence were confirmed by the correlation across item positions in the sequence (Figure 4D). A simple mechanism that could explain this confidence-frequency relation will be presented below.

Before doing so, one might underline a striking similarity between the present finding that confidence is blind to selection delays in the Attentional blink paradigm and a previous finding that introspective response times are blind to delays induced by the psychological refractory period – or PRP – (Corallo, Sackur, Dehaene, & Sigman, 2008; Marti, Sackur, Sigman, & Dehaene, 2010). In this paradigm, two tasks have to be conducted on stimuli presented in short succession in time, and it has been shown that the decision process for the second task is postponed until the decision process for the first task has been completed. Interestingly however, introspective estimates of response times are blind to this delay induced by the serial central bottleneck. It has been suggested that the Attentional blink and PRP paradigm involve a similar central bottleneck (Marti, Sigman, & Dehaene, 2012; Wong, 2002). Indeed, introspective measures of performance (respectively, subjective estimates of accuracy, i.e. confidence, and subjective estimates of response times) appear to be oblivious to the delays induced by this central bottleneck in both paradigms. To expand this research, future work might investigate whether introspection is blind to central delays in different paradigms, or to other constraints of central processing stages (e.g. the discrete/symbolic nature of information processing at central stages, see de Gardelle, Charles, & Kouider, 2011; de Gardelle, Kouider, & Sackur, 2010).

### Confidence follows report frequency: a candidate mechanism

Confidence can build on more than the decision evidence. Studies on confidence and the first-order decision (accuracy) have resulted in two main observations: most of the time, confidence successfully tracks accuracy, but systematic dissociations exist (e.g., Graziano & Sigman, 2009; Lau & Passingham, 2006; Peters et al., 2017; Rahnev et al., 2011; Rahnev et al., 2012). Those dissociations have led to reevaluate the historical view that both first-order decisions and confidence judgments are made solely from the same evidence signal (Mamassian, 2016). In this context, a third main finding of the present work is that confidence generally follows report frequency, across positions of the items in the sequence, irrespectively of whether T1 or T2 is considered, and irrespectively of the T1-T2 lag or the delays induced by the blink. In other words, irrespectively of whether attention is oriented or reoriented to a second point in time. This simple relationship between confidence and report frequency can explain many aspects of our data, such as (i) the error-based metacognition (how confidence was sensitive to objective errors) for T1, (ii) confidence being blind to delays induced by the Attentional blink, and (iii) the degradation of error-based metacognition for T2 during the Attentional blink.

Interestingly, the finding that confidence is related specifically to report frequency is also well accounted for by a simple addition to a classic model of item selection during RSVP tasks. In this classic model, items encoded within the perceptual system receive an attentional boost, which is smoothly distributed in time over several items (see e.g. Figure 15 in Reeves & Sperling, 1986). At the end of the sequence, the reported item is the one with the highest evidence (which is the product of encoding strength and attentional boost). In addition, random perturbations (noise) on the final evidence levels, and/or constraints at the encoding or attentional stage (e.g. delays imposed by the selection of a first target), may move the peak evidence away from the correct target, allowing for random errors and systematic delays in item selection (Vul, Hanus, & Kanwisher, 2009).

This model could also predict our results regarding confidence, if we additionally assume that confidence corresponds to the amount of evidence for the selected item – or first-order decision. In this version of the model, systematic delays in attentional boost would shift both item selection and confidence in the exact same manner within a sequence, such that confidence would be blind to selection delays. Furthermore, random perturbation on evidence levels would induce a correlation between report frequency and confidence. To understand why, one might consider T1 data (Figure 2, A and B): reports frequencies indicate that the evidence peak for T1 selection is often correctly located at the target position. Noise added to the evidence of each item could however shift the peak evidence to a different position, but positions further away from the target would still be selected less often, and when selected, their evidence would still be attenuated relative to positions closer to the target. What follows is a correlation between report frequency and confidence across positions, as was found in our data.

## Conclusion

The strong correlation between frequency of reports and confidence during temporal selection (T1), which holds when attention has to reorient to a second point in time (T2), suggests that first decision and confidence are mostly sharing the same evidence signal during temporal orienting of attention. This tight coupling might prevent confidence from accessing delays in selection induced by the Attentional blink, as shown in the present work. In addition to this confidence-frequency relation, lies a more general heuristic that applies a confidence cost to items selected at early stages of attentional reorienting, where a second target benefits from a prior attentional episode (Lag-1 under-confidence). These multiple phenomena highlight a need for further work on the hypothesis that confidence and temporal attention might operate at different time scales, a difference which could reflect late heuristic biases, rather than segregated processes for decision and confidence evidence.

## Supporting information

