## Supplementary material for "When temporal attention lacks confidence"

Samuel Recht[1]*, Pascal Mamassian[1], Vincent de Gardelle[2]

[1] Laboratoire des systèmes perceptifs, Département d'études cognitives, École normale supérieure, PSL University, CNRS, 75005 Paris, France

[2] CNRS and Paris School of Economics, Paris, France

### Index

### Experiment 1

#### Order reversals between T1 and T2

Order reversals occur at lag-1 when participants report both T1 and T2 but in the reverse order. In our data, order reversals occurred on average in 7.68% (SE ± 4.71%) of lag-1 trials. For comparison, correct report of both T1 and T2 in the correct order occurred in 12% (SE ± 5.77%) of lag-1 trials. To evaluate whether participants were aware of such reversals, the confidence between trials in which both T1 and T2 were correctly reported was compared to the confidence in reversed trials. One participant was discarded from this analysis due to no order reversal trial. No difference in confidence was found between these two types of trials, neither for T1 ($t(30)=1.07$, $p=0.29$) nor for T2 ($t(30)=1.20$, $p=0.24$). Thus, it seems that participants were not specifically aware of the occurrence or non-occurrence of a reversal on a trial-by-trial basis. However, it is still possible that participants could be aware of the possibility of order reversals at lag-1 relative to longer lags, and that being aware of this possibility would be responsible for the lag-1 under-confidence.

#### Position-based metacognition

In a finer analysis, we tested whether participants' confidence could discriminate between different errors across different serial positions, not just between correct and incorrect responses. Excluding correct T1 responses, we found that a regression model with position effect and lag outperformed the null model without the position for predicting confidence ($\chi^2(3)=101.2$, $p_{RAND}<0.001$), with no significant interaction between position and lag ($\chi^2(12)=7.99$, $p_{RAND}=0.78$). Participants are thus sensitive to the difference between various position errors, even if this distinction is irrelevant to succeed in the present task.

### Experiment 2: a replication with lowered metacognitive load

#### Material & methods

##### Participants

35 adult volunteers were recruited from the Laboratoire d'Economie Expérimentale de Paris (LEEP) pool of participants (M ± SD = 24.5 ± 3.06 years old). They all provided informed written consent prior to the experiment. One observer was discarded for not finishing the experimental session, and 6 participants were removed following extremely small accuracy rate for target 1 or 2 (exclusion criterion: <10% accuracy), leaving 29 participants for analysis. Observers were paid a base sum (10 EUR) plus a bonus depending on their performance in the task (up to 10 EUR in addition). The average payoff was 14.89 EUR (± 2.09) for a single 1.5 hours session. The experimental procedure received approval from the Paris School of Economics (PSE) ethics review board and adhered to the principles of the Declaration of Helsinki.

##### Apparatus and stimuli

Identical apparatus, stimuli and parameters were used for both experiments. The only difference being that for Experiment 2, confidence judgment was required only for T1 on half of the 500 trials, and only for T2 on the other half. Participants were divided into two groups to control for possible order effect. Participants were left uninformed that they will have to estimate their confidence for the other target until the end of the first half of the experiment.

#### Analysis

For the following analyses, trials were grouped by confidence probe: one group of trials for T1 confidence (250 trials per participant) and one group of trials for T2 confidence (250 trials). Therefore, even when accuracy only was considered, the average concerns the subset of trials related to the target where confidence judgment was requested.

### Results

#### T1: distribution of reports

The results from Experiment 1 were successfully replicated, with a significant effect of lag on accuracy ($F(3.4, 5.134)=9.1$, $MSE=0.005$, $p<0.001$) and confidence ($F(1.65,46.17)=17.5$, $MSE=0.08$, $p<0.001$). We found that letters presented just before or just after the target were reported on 19% of the trials (18% in Exp. 1), which exceeded the guess rate of 1/26 that is about 4% (mean corrected for guess rate: 0.15, 95% CI=[0.13 0.17]; $t(28)=14.7$ $p<0.001$).

To quantify how report frequency depended on serial position, we focused on serial positions from 2 items before to 2 items after T1 (included) and tested how report frequency can be predicted from the lag, the position and their interaction as fixed effects. Including item position as a predictor outperformed a model without the position effect ($\chi^2(4)=565.0$, $p_{RAND}<0.01$). Including the interaction between lag and position had not improved the model over a model without the interaction ($\chi^2(16)=21.6$, $p_{RAND}=0.36$), contrary to Exp. 1.

#### T1: Confidence follows report frequency

Similarly to Exp. 1, confidence was affected by item position ($\chi^2(4)=94.03$, $p_{RAND}<0.01$). Including the interaction between lag and position however did not improve the model ($\chi^2(16)=26.0$, $p_{RAND}=0.16$). Given that for T1 data, both report frequency (figure S1A) and confidence (figure S1B) were affected by position in similar manners, we directly evaluated the correlation between confidence and report frequency. To do so, for each participant we averaged confidence over lags, and correlated this average confidence to the report frequency across 5 report positions centered on target (including target's true position). The mean r coefficient was 0.71 across participants (95% CI=[0.59 0.83]; $t(27)=11.9$, $p_{RAND}<0.001$), replicating Exp. 1.

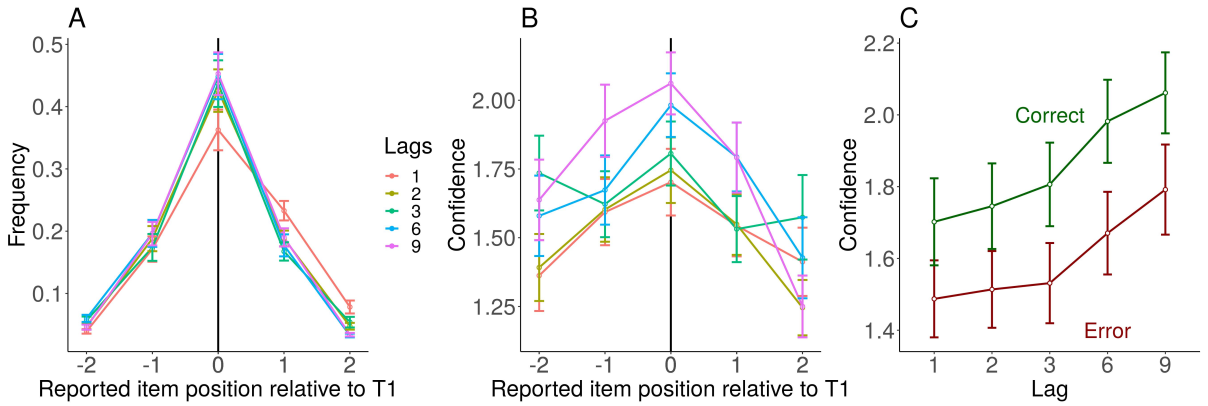

**Figure S1** | **Reports and confidence about T1** (A) The frequency of report for item around target true position. (B) The corresponding average confidence per position. (C) The correct and error trials average confidence level, which provides an estimate of objective, error-based metacognition. Error bars represent standard error of the mean across participants.

#### T1: Error-based metacognition

Overall, T1 targets in Exp. 2 – as for Exp. 1 - were identified correctly 43% of the time. A main effect of trial type (error versus correct trial) was found ($F(1,28)=39.0$, $MSE=0.13$, $p<0.001$) but no interaction between lag and trial type ($F(3.598,100.737)=1.2$, $MSE=0.02$, $p=0.31$), confirming that participants had stable error-based metacognition for T1. Participants therefore gave higher confidence to correct than to incorrect T1 responses.

#### T2: Confidence tracks the attentional blink

Overall, 19% of T2 reports were correct when T1 was correctly reported. Figure S2A shows T2 accuracy and confidence for the different T1-T2 lags. As expected, the accuracy of T2 reports (i.e. in green) was affected by the lag between T1 and T2 ($F(2.9,81.13)=41.4$, $MSE=0.03$, $p<0.001$). In particular, the drop for lag 2 and lag 3 relative to longer lags (2-3 vs. 6-9: $T(28)=7$, $p<0.001$) indicated a classical Attentional Blink effect. Confidence was also affected by lag ($F(2.59,72.42) = 37.2$, $MSE=0.10$, $p<0.001$) and dropped for lags 2-3 relative to longer lags (2-3 vs. 6-9: $T(28)=0$, $p<0.001$), paralleling the drop observed for accuracy. Thus, participants seem able to acknowledge the drop of performance during the attentional blink that occurs at lags 2-3, in a similar manner as for Exp. 1.

Participants' confidence, similar to Exp. 1, seemed blind to lag-1 sparing. Indeed, the lag-1 sparing effect was also found in our data: T2 accuracy was spared when T2 was presented immediately after T1. Accuracy at lag-1 was much higher than during the blink period (1 vs. 2-3: $T(28)=378$, $p<0.001$) and was in fact was indistinguishable from accuracy at long lags (1 vs. 6-9: $T(28)=238$, $p=0.67$). By contrast, confidence was as low at lag-1 as it was for lag 2-3 ($T(28)=160$, $p=0.70$) and much lower than confidence at long lags (1 vs. 6-9: $T(28)=9$, $p<0.001$). All these results were fully coherent with what was found in Exp. 1.

**T2: underconfidence during Lag-1 sparing**

As in Exp. 1, a normalization procedure was applied to further quantify "lag-1 under-confidence". Figure S2B shows confidence and accuracy for Exp. 2 at lag-1, in this lag-3-to-9 space, where lag-3 and lag-9 have (0,0) and (1,1) coordinates, respectively. Most participants are located below the diagonal, suggesting that they are less confident at lag-1 than what would be expected given their accuracy level at lag-1. This lag-1 underconfidence, computed as the average difference between predicted and observed lag-1 confidence, was significant at the group level ($M=0.51$, 95% CI=[0.30 0.72]; $t(27)=5.0$, $p<0.001$). These results suggest no strong influence of metacognitive load on the level of underconfidence observed during lag-1 sparing: probing confidence only for T2 had not altered the original pattern.

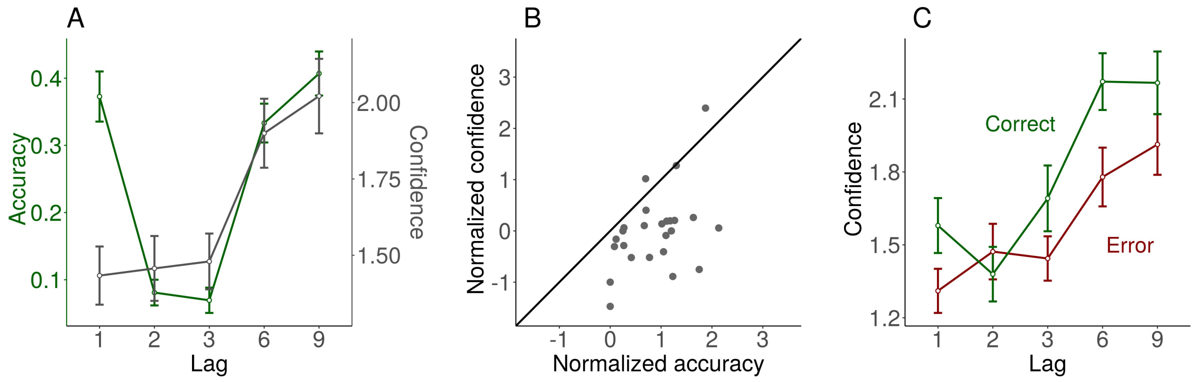

**Figure S2 | Attentional Blink and early confidence bias under lowered metacognitive load**. (A) T2 average accuracy (in green) and confidence (in grey) as a function of the lag between T1 and T2. (B) The systematic under-confidence occurring when attention reorients 80ms after the first target (Lag-1), by representing the accuracy and confidence of lag-1 as a point in the space inferred from the covariation of confidence and accuracy from lag-3 to lag-9. Each point is a participant. (C) The average confidence level for correct T2 reports and errors (error-based metacognition), for each lag. Metacognition is conserved at lag-1 despite low confidence ratings. All results are similar to the results in Experiment 1.

**Order reversal between T1 and T2**

In Exp.2, order reversals occurred on average in 5.2% (SE ± 4.1%) of lag-1 trials. For comparison, correct report of both T1 and T2 in the correct order occurred in 13.2% (SE ± 6.3%) of lag-1 trials. To evaluate whether participants were aware of such reversals, the confidence between trials in which both T1 and T2 were correctly reported was compared to the confidence in reversed trials. Seven participants were discarded from the later analysis due to no order reversal trial for the T1 confidence block, and two participants were discarded for the T2 confidence block. No difference in confidence was found between these two types of trials for T1 ($t(21)=0.4$, $p=0.70$), but a near significant difference was found for T2 ($t(26)=2.2$, $p=0.04$). Thus, it seems that when metacognitive load is reduced, participants were aware of the occurrence or non-occurrence of a reversal on a trial-by-trial basis, but only when confidence was requested for T2.

**T2: Error-based metacognition**

Because some participants had no correct answers during the AB, only half of participants were considered here (N=13). As can be seen from Figure S2C, participants overall expressed higher confidence when they were correct relative to their errors ($F(1,13)=16.5$, $MSE=0.10$, $p=0.001$). This error-based metacognition interacted with the lag ($F(3.13,40.67)= 21.8$ $MSE=0.16$, $p<0.001$). Post-hoc Bonferroni-corrected tests showed that the difference in confidence between correct T2 reports and errors was significant for lag-1 and lag-9 ($p<0.002$) but not for other lags. The ability to detect objective errors was therefore, as in Exp.1, diminished during the AB period, and it did not disappear at lag-1, despite the low level of confidence.

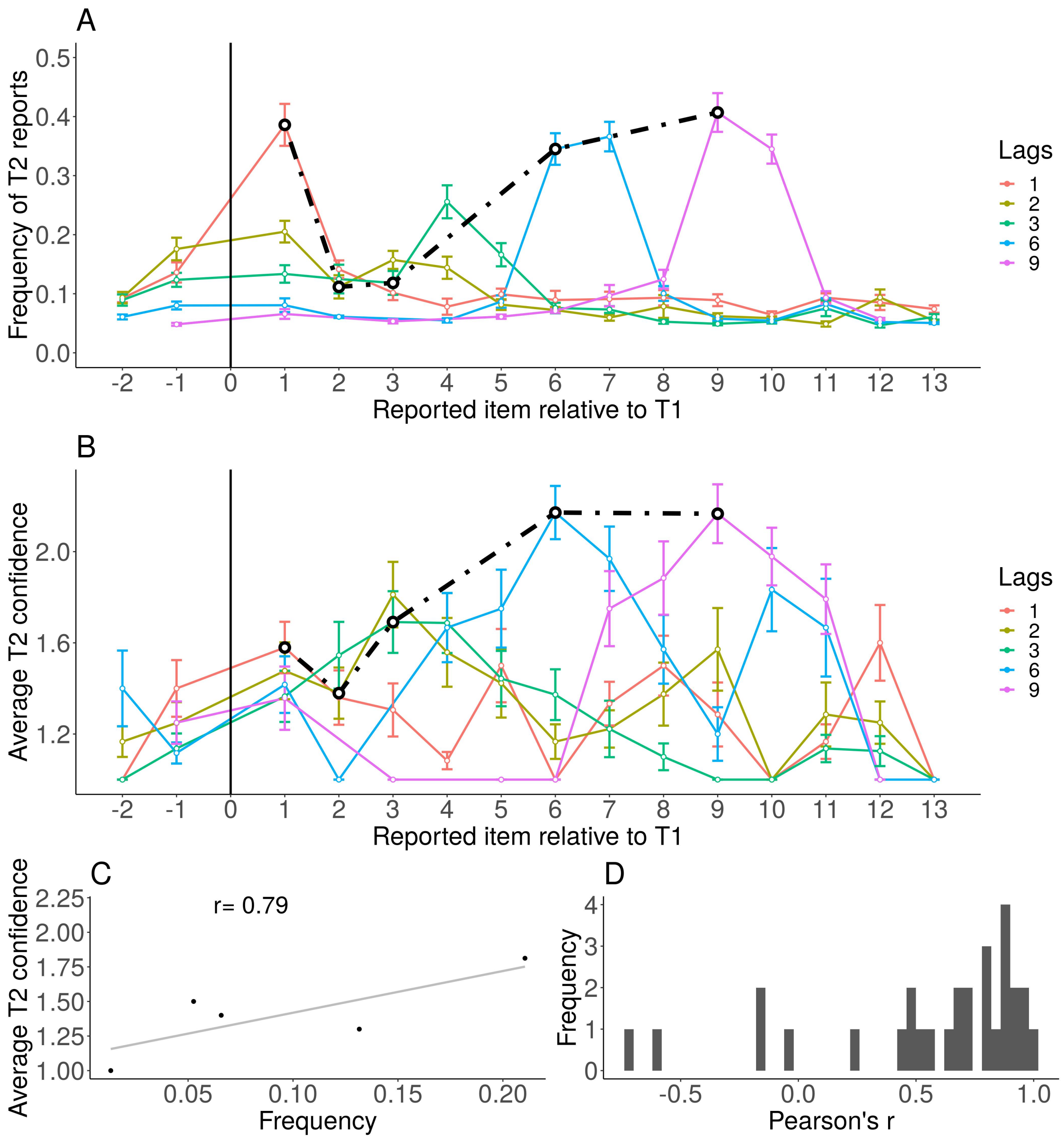

**Figure S3** | **Reports and confidence about T2**. (A) The frequency of T2 reports as a function of the position of the reported item relative to T1, for each lag. Note that T1 position has no value, given that only trials in which T1 is correctly reported were considered here (hence T2 reports cannot correspond to T1 position). The black line connects the points corresponding to accurate T2 reports. (B) Confidence of the T2 reports, as a function of the position of the reported item relative to T1, for each lag. The black line connects the points corresponding to accurate T2 reports. Error bars represent standard error of the mean across participants. (C) Regression between frequency and confidence with 5 positions centered on T2, collapsed across lags, for a representative participant.

(D) Histogram of the correlation coefficients for all the participants, the correlation is not significant using permutation test. Please refer to the online version of the paper for colors.

**T2: Delay in temporal selection and confidence**

As for Exp. 1, items appearing just before or just after the T2 were more likely to be reported than chance (17%, with a 95% CI=[0.15 0.18]; vs. chance level at 4%: $t(28)=19.0$, $p<0.001$). Hence, errors were not random guesses but samples that are close to the actual T2 target. Replicating Exp. 1, selection appears to be systematically too late for lags 3, 6 and 9.

A model comparison approach confirmed that including the lag as a predictor for the center of mass significantly outperformed the null model ($\chi^2(4)=18.4$, $p_{RAND}=0.001$). Furthermore, a model including confidence, lag and their interaction as fixed effects outperformed a model without the interaction term ($\chi^2(8)=19.0$, $p_{RAND}<0.05$), which in turn outperformed a model without the main effects of confidence and lag ($\chi^2(2)=15.7$, $p_{RAND}<0.01$). When looking at each lag separately, we found that the effect of confidence on the center of mass was only significant for lag-3 after Bonferroni correction ($\chi^2(4)=15.4$, $p_{RAND}<0.01$) but not for the other lags ($p>0.1$). For lag-3, reports given with higher confidence are also associated with longer delays than reports given with low confidence (figure S4 right panel). In other words, confidence is oblivious to the delays induced by the AB and biased towards items selected later. A reduced metacognitive load in Exp. 2 did not enhance delay introspection, on the contrary.

**T2: Confidence and frequency**

The similarity between confidence and report frequency was tested by looking at their correlation across lags for 5 positions centered on T2, but contrary to T1, the correlation was not reaching significance (Mean r coefficient: 0.55, 95% CI=[0.38 0.73]; $t(31)=6.5$, $p_{RAND}=0.06$), as shown on figure 4D. Figure 4C plots the regression on one representative participant for illustrative purpose. The smaller correlation found in Exp. 2 compared to Exp. 1 might be the result of the reduced number of samples (half of Exp.1 samples for T2 confidence).

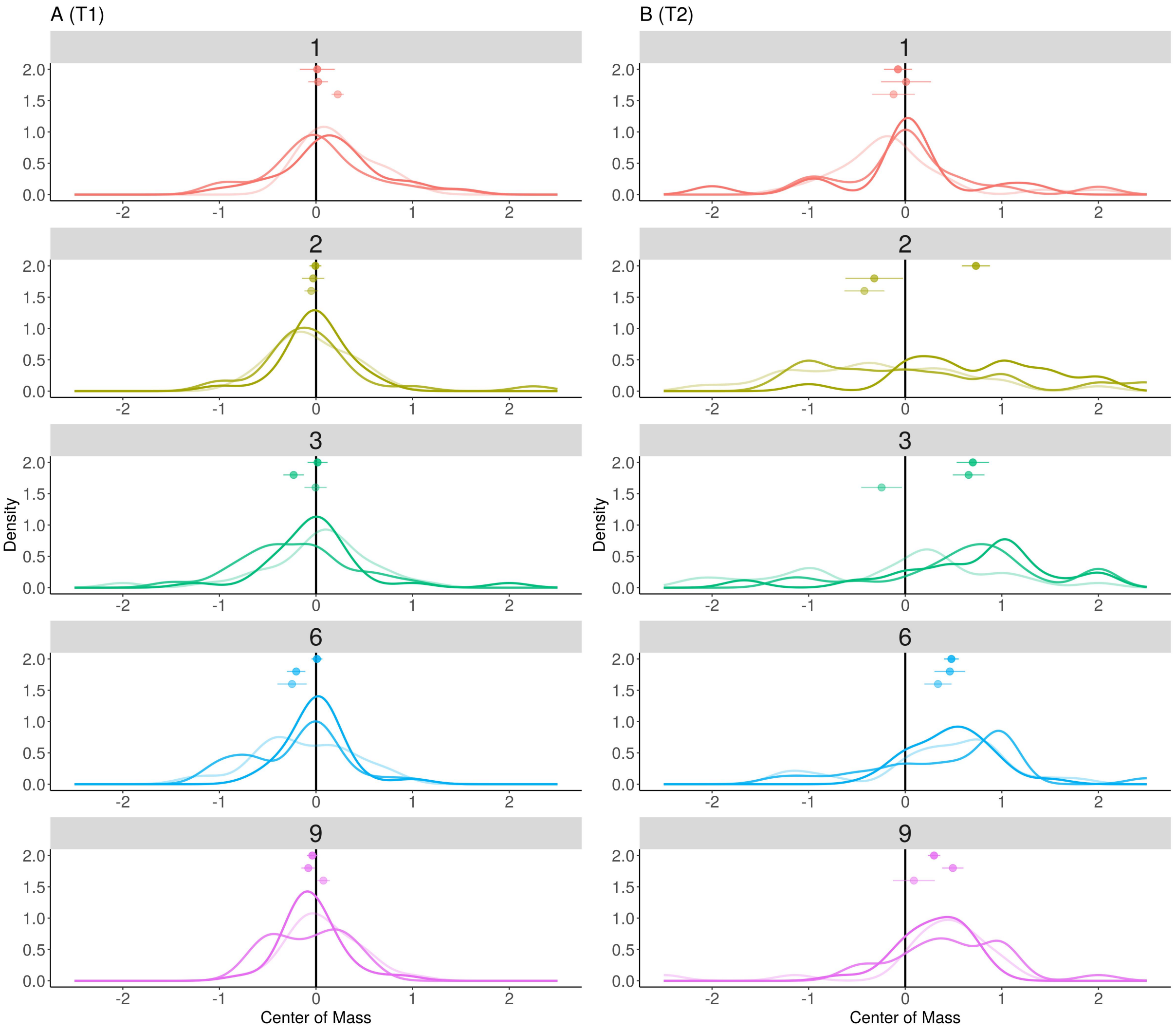

**Figure S4 │ Center of mass analyses.** Each panel represents for one lag the distribution of the center of mass across participants, relative to the T1 or T2 positions. The 3 confidence levels are plotted separately for each lag, with more transparent lines for lower confidence. Note that at lag-3, higher T2 confidence is associated with longer delays, suggesting that confidence does not accurately track the delay of temporal attention. Dots and error bars represent the mean and standard error of the mean for each distribution. Densities were estimated using Gaussian kernels (sd: 0.2).
